# Addressing confounding artifacts in reconstruction of gene co-expression networks

**DOI:** 10.1101/202903

**Authors:** Princy Parsana, Claire Ruberman, Andrew E. Jaffe, Michael C. Schatz, Alexis Battle, Jeffrey T. Leek

**Author notes:** Equal Contribution (Princy Parsana, Claire Ruberman). Corresponding authors: Alexis Battle -; Jeffery T. Leek.

## Abstract

**Background:** Gene co-expression networks capture diverse biological relationships between genes, and are important tools in predicting gene function and understanding disease mechanisms. Functional interactions between genes have not been fully characterized for most organisms, and therefore reconstruction of gene co-expression networks has been of common interest in a variety of settings. However, methods routinely used for reconstruction of gene co-expression networks do not account for confounding artifacts known to affect high dimensional gene expression measurements.

**Results:** In this study, we show that artifacts such as batch effects in gene expression data confound commonly used network reconstruction algorithms. Both theoretically and empirically, we demonstrate that removing the effects of top principal components from gene expression measurements prior to network inference can reduce false discoveries, especially when well annotated technical covariates are not available. Using expression data from the GTEx project in multiple tissues and hundreds of individuals, we show that this latent factor residualization approach often reduces false discoveries in the reconstructed networks.

**Conclusion:** Network reconstruction is susceptible to confounders that affect measurements of gene expression. Even controlling for major individual known technical covariates fails to fully eliminate confounding variation from the data. In studies where a wide range of annotated technical factors are measured and available, correcting gene expression data with multiple covariates can also improve network reconstruction, but such extensive annotations are not always available. Our study shows that principal component correction, which does not depend on study design or annotation of all relevant confounders, removes patterns of artifactual variation and improves network reconstruction in both simulated data, and gene expression data from GTEx project. We have implemented our PC correction approach in the Bioconductor package *sva* which can be used prior to network reconstruction with a range of methods.

## Background

Most cellular processes are carried out by groups of genes that function together, often supported by coordinated transcriptional regulation. Based on this, gene co-expression networks seek to identify transcriptional patterns that are indicative of functional interactions and regulatory relationships between genes[1–3], which are not yet fully characterized for most species, tissues, and disease-relevant contexts. Therefore reconstructing co-expression networks from high-throughput measurements is of common interest. However, accurate reconstruction of such networks remains a challenging problem.

Co-expression networks can be encoded as undirected graphs where nodes represent genes, and edges between nodes indicates correlated expression between genes. Edges indicate a potentially functional relationship between the genes such as a transcription factor and its gene targets, pairs of genes within a shared pathway, or genes within a protein complex. Widely used network learning methods such as Weighted Gene Co-expression Network Analysis (WGCNA)[4]and graphical lasso[5]are based on pairwise associations between genes[4,5].Some specialized methods for reconstruction of co-expression networks consider confounding signals within their model[6,7],however most commonly used methods do not directly account for artifacts such as batch effects, sequencing artifacts, and unwanted biological effects. It has been shown that these artifacts can affect inference of co-expression in high-dimensional gene expression data [8].Both biological and technical artifacts are known to influence expression measurements of gene expression data, sometimes substantially, introducing spurious signals including false correlations between genes which are not reflective of a functional relationship(Chen et al. 2011; Leek and Storey 2007).Some of the major confounders known to impact RNA-seq measurements include RNA quality (like RNA integrity number, RIN), RNA extraction and library preparation batches, sequencing library size, mapping artifacts, and GC bias[11–14].Biological confounders such as age, sex, ancestry, cell type heterogeneity and others can lead to apparent co-expression of genes with no direct functional interaction. Additionally when gene expression data is collected from organ donors, covariates such as hardy scale, ICD-10 code for cause of death, core body temperature, and other subject specific attributes also significantly affect gene expression measurements[14].However, while it has been long known that known and hidden artifacts introduce erroneous correlations in the gene expression measurements, many studies yet do not employ any form of correction of gene expression data prior to network reconstruction (Supplementary Table 1). Correlations introduced by these confounders are often inferred as relationships between genes, leading to inaccurate network structure and erroneous conclusions in downstream analyses[6,7,9,15,16]. Therefore, it is critical to correct gene expression data for unwanted biological and technical variation without eliminating signal of interest before applying standard network learning methods.

In this study, we provide a framework for data correction leveraging the structure of scale-free networks. It has been shown that real world networks including co-expression networks often have scale-free topology, i.e. the node degree distribution of these networks follow a power law[17–19]. These networks are characterized by a small number of influential hub nodes that link to the remainder of the lower degree nodes. This makes scale-free networks more stable and more robust to random perturbations[20–25].Several studies have employed the assumption of scale-free topology to infer high-dimensional gene co-expression and splicing networks[4,26].

Here, we show that for scale-free networks, principal components of a gene expression matrix can consistently identify components that reflect artifacts in the data rather than network relationships. The number of principal components can be estimated in multiple ways, here we use a permutation based scheme[27]for estimating the number of principal components as implemented in the sva package[28].We prove that under a scale free network that these principal components can then be removed without affecting the signal arising from the true gene associations, enabling improved network reconstruction. While not every biologically meaningful network structure will satisfy these assumptions, this framework provides a useful and simple approach for removing major confounding signals while preserving many biologically relevant relationships.

## Results

### Principal components of gene expression matrix reflect artifacts in the data

It is now well-known that technical confounders can have broad impact on genomic measurements[10,15,29,30]. Since the signal from unmeasured or unmodeled artifacts often accounts for a large fraction of variance in genomic data, latent variable methods such as principal components analysis, factor analysis or surrogate variable analysis can be used to detect and estimate these artifacts, which can then be adjusted for in downstream analyses [31,32]. These approaches have been successfully applied to gene expression data with methods such as RUV[33],SVA[10],and PEER[29].Similar approach have also been applied to identify and control for confounders in genetic association studies with methods like EIGENSTRAT[30,34].

In differential expression analysis, unmeasured confounders may lead to pervasive spurious signal across many features (genes) simultaneously[10,11,33].However, it is also possible that a strong biological signal of interest - for example cancer versus control status - may lead to biologically meaningful changes in gene expression for many genes. When we are interested in differential expression analysis with a single or small set of desired biological factors, supervised latent variable methods such as SVA, RUV, and PEER can adjust the estimation of latent confounders to ensure that relevant biological signal is retained[10,15,29]. In contrast, in genome-wide association studies investigating association between genotype and complex traits, adjustment for latent factors has been used successfully without supervision, because broad correlation between genotypes generally reflects population structure rather than desired functional biological signal of interest. Unfortunately, co-expression analysis is more complicated than either differential expression or GWAS because the potential for arbitrary unknown confounders remains, and can affect sets of genes in ways that resemble co-expression. When commonly measured covariates such as RIN, batch, exonic rate, gene GC%, and others are available their effect from gene expression data can be regressed out using a linear model. Unfortunately more detailed annotations are generally not available for a wide range of confounding variables such as post mortem interval, environmental factors, or detailed technical records. Furthermore, in many cases when densely annotated data is available for technical and biological artifacts, they are not complete, i.e. some covariates are not recorded for all observations thereby making correction more challenging.

Here we show mathematically, through simulation, and through real data examples that reconstruction of gene co-expression networks resembles the parameter regime of genetic association studies, i.e. we expect that most real co-expression networks are sparse which means that there are few true biological connections between genes. When such networks satisfy the small world property, the signals from the network will not be sufficiently broad across genes to influence the latent variable estimates from PCA. Thus the principal components will primarily estimate the latent confounders which can then be regressed from the expression data before network reconstruction is performed on subsequent residuals (**Supplementary Note 1**).

### Confounders can introduce spurious relationships in co-expression networks - a simulated example

Using a small simulated example, we illustrate how confounders in gene expression data can impact reconstruction of co-expression networks and how this can be corrected (**Figure 1**). We used three versions of data -- (a) simulated gene expression data from a sparse network with no batch effects (**Figure 1a**); (b) simulated data with a batch effect added (**Figure 1d**); and (c) confounded data from (b) corrected by regressing out the top principal component (**Figure 1g**). With each version of the data, we first computed empirical correlation matrix (**Figure 1b,e,h**). Next, we reconstructed co-expression networks using graphical lasso[5,35].Confounders in the simulated data that affected genes 2 through 6 was evident through the block pattern in the data matrix (**Figure 1d**). Likewise a large block of high (spurious) correlation between the same genes was observed in the empirical correlation matrix (**Figure 1e**). Moreover all genes that were affected by confounders were connected to each other in the inferred network while two true dependencies *E(3,1)*and *E(4,7)*were lost (**Figure 1f**).

**Figure 1.**
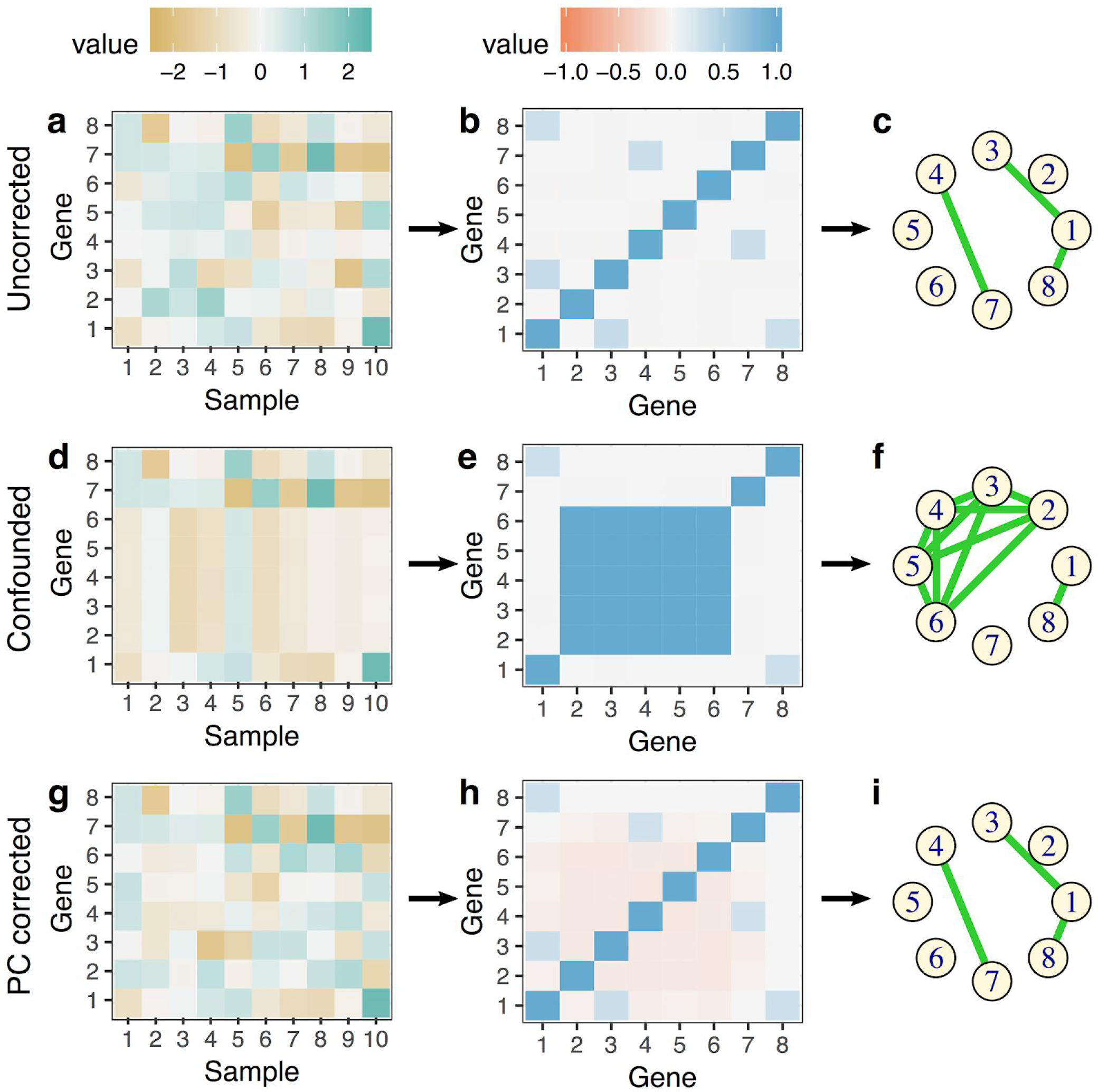
A simulation example. This simulation example shows reconstruction of gene co-expression networks is affected by confounders. True underlying network structure can be reconstructed after principal component correction of gene expression data as described in the paper.

Next we corrected the confounded data using a linear model with the top principal component in the confounded data as an explanatory variable. We then reconstructed the network using the residuals from this regression (**Methods**) The structure and pattern of heatmap and correlation matrices of the corrected data resembled the original simulated data with some additional negative correlation (**Figure 1a-b** **and g-h**). Additionally, graphical lasso correctly estimated the network structure obtained from corrected data, which was same as the true network structure that was obtained from the original simulated data (**Figure 1c** **and i**).

### Principal component correction improves reconstruction of co-expression networks in GTEx tissues

To demonstrate the effect of latent confounders and principal component correction on reconstruction of co-expression networks from real large-scale human gene expression measurements, we applied our method to RNA-seq data from the Genotype Tissue Expression (GTEx) project v6p release. We considered data from eight diverse tissues containing between 304 and 430 samples each: Subcutaneous adipose, Lung, Skeletal muscle, Thyroid, Whole blood, Tibial artery, Tibial nerve and Sun-exposed skin. We used two popular methods for network reconstruction: (a) weighted gene co-expression network analysis[4,36],and (b) graphical lasso[5,35]. Since the true underlying co-expression network structure is not known, we assessed the networks using genes annotated to function in the same pathways, and transcription factor with known target genes as the ground truth (**Methods**).

We obtained the most recent version (2016) of curated gene sets annotated as canonical pathways on MsigDB[37] available on the Enrichr library, containing information from KEGG, Reactome, Biocarta and Pathway Interaction Database. False discovery rate was used to evaluate and compare the performance of co-expression networks inferred from data after correcting for known or latent confounders. We evaluated networks inferred using a) uncorrected expression data, the residuals after regressing out b) RNA integrity number (RIN), c) exonic rate - a mapping covariate that corresponds to fraction of reads mapped to exons, d) sample specific estimate of GC bias, all shown to be common confounders in mRNA gene expression data, and e) residuals from multiple regression model using covariates that explained at least 1 percent of expression variance (adjusted *R*^2^ ≥ 0.01, **Supplementary Table 4)** [12,38–41].

WGCNA identifies groups of genes that form coexpressed modules based on a power transform of the pearson correlation coefficient for all pairs of genes[4].For each tissue we inferred weighted, signed co-expression networks using the most variable 5000 genes (**Methods**). Co-expression gene modules were identified based on fully-connected sub-graphs of the network. We find that for most tissues networks obtained from data corrected for latent confounders showed fewer false discoveries compared to those obtained from uncorrected data, or from correcting for individual covariates including RIN, exonic rate (a quality metric from RNA-seq mapping), or sample-specific GC bias (**Figure 2**, **Supplementary Figure 1, 3, 6**). Improved performance of networks obtained from PC corrected data was more evident in whole blood, skeletal muscle, tibial artery, tibial nerve, subcutaneous adipose and thyroid. But for some tissues such as lung, PC correction only contributes to moderate improvement on false discovery rates in the reconstructed networks. It is possible that in these cases, the networks may violate the small world assumption. We also observed that correcting gene expression data with multiple technical covariates (approximately 9 - 17 were used per tissue) sometimes improved reconstruction of co-expression networks obtained by WGCNA(**Figure 2**, **Supplementary Figure 1**). We also inferred signed WGCNA networks with varying values of the power transform parameter, and unsigned co-expression networks at varying cut-heights of the hierarchical dendrogram. With varying power transform we found that PC correction improved reconstruction of WGCNA networks in all tissues (**Supplementary Figure 15**). Most tissues except subcutaneous adipose showed reduced false discoveries in unsigned WGCNA networks. (**Supplementary Figure 14**).

**Figure 2.**
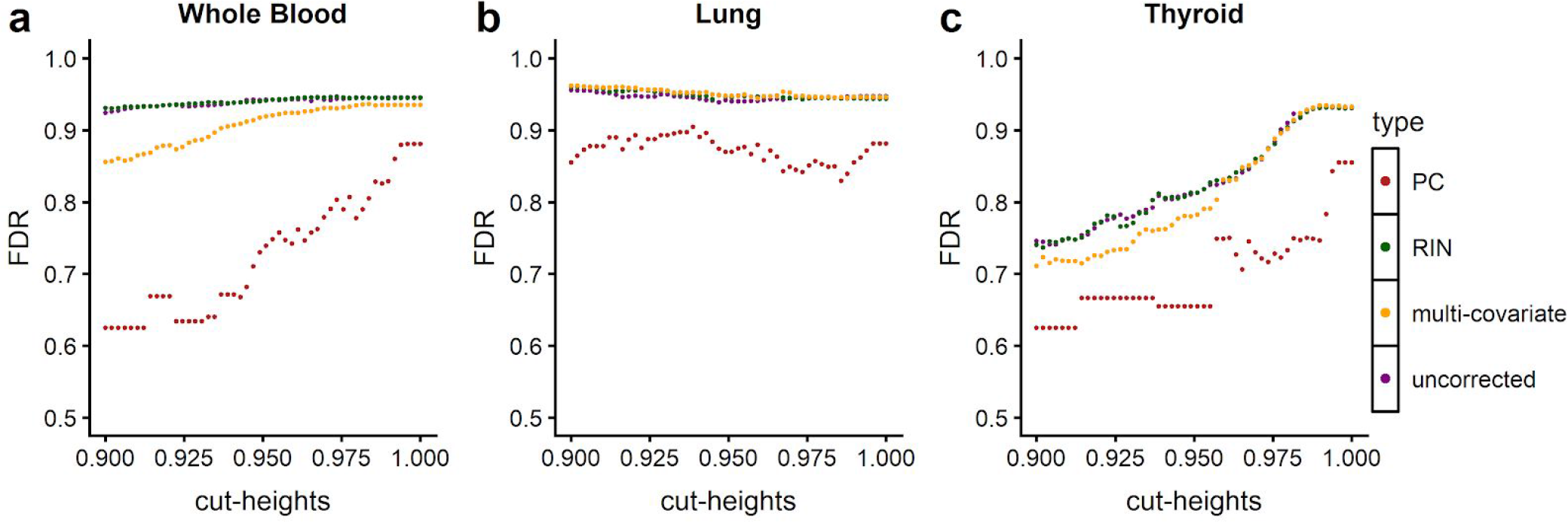
False discovery rate of WGCNA modules based on canonical pathways. FDR of WGCNA networks obtained at a varying cut-heights. Each point corresponds to FDR of the network obtained at a specific cut-height. Each color represents networks reconstructed with a specific correction approach

Similarly we examined the effect of confounders on networks reconstructed with graphical lasso using the same 5000 most variable expressed genes in each tissue. To test the effect of sparsity we varied the penalty parameter in graphical lasso (λ=[0.3,1.0]). For each tissue, using the non-zero entries in the estimated precision matrix as the edges of the graph, we computed false discoveries for the inferred networks (**Methods**).

We found that glasso networks estimated with principal component corrected data showed fewer false discoveries compared to the networks estimated with uncorrected, RIN corrected or multiple covariates corrected data (**Figure 3**, **Supplementary figure 2**). We observed that in general improved performance on false discoveries in PC corrected networks was better in whole blood, skeletal muscle, tibial artery and tibial nerve. Jointly correcting the gene expression data for multiple technical covariates that affect expression measurements, improved reconstruction of co-expression networks with graphical lasso in some tissues such as whole blood, thyroid, and tibial artery, while it showed little to no improvement over uncorrected data in lung, muscle, tibial nerve, and sun exposed skin(**Figure 3**, **Supplementary figure 2**). However we observed that across all tissues PC correction still shows fewer false discoveries compared to multiple technical covariate based correction. Similar to WGCNA we observed that there was no visible improvement in network reconstruction between using uncorrected data and residuals from RIN or exonic rate; thereby suggesting that RIN, exonic rate or GC bias individually is not a sufficient alternative for the wide range of confounding variation found in gene expression data (**Figure 3**, **Supplementary Figure 4, 7, 9, 11**).

**Figure 3.**
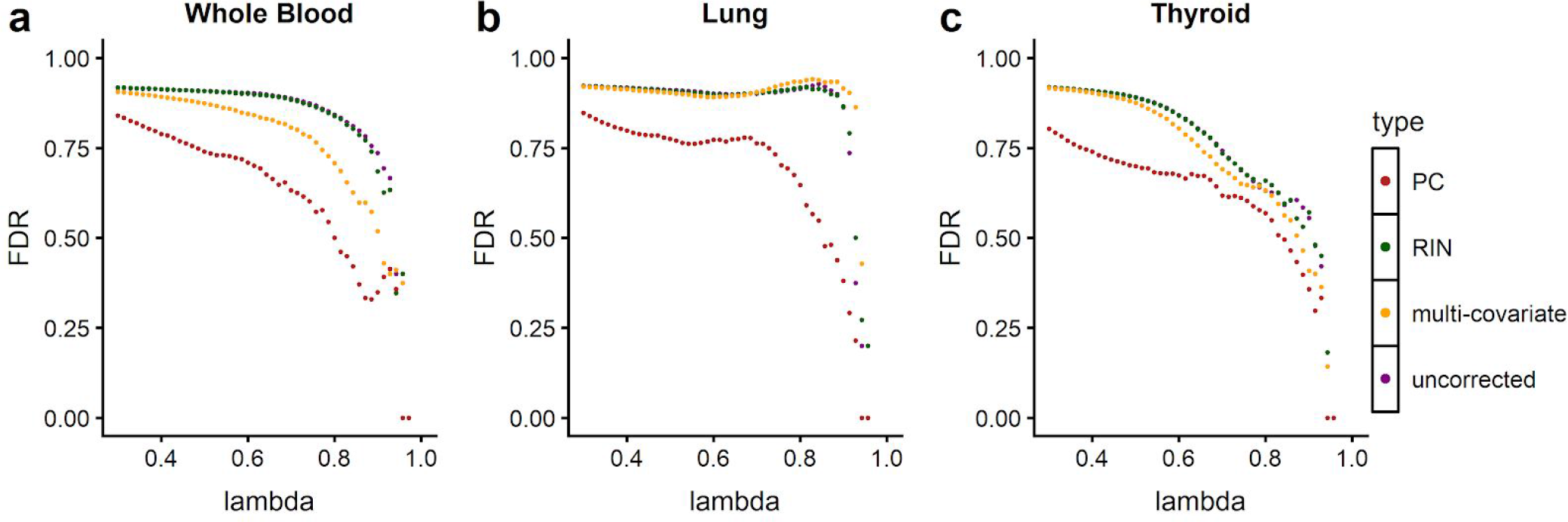
False discovery rate of networks inferred with graphical lasso based on canonical pathways. Each point in the figure corresponds to false discovery rates of networks obtained at a specific L1 penalty parameter value. Each color represents networks reconstructed with a specific correction approach - uncorrected, multi-covariate, RIN, and PC corrected

We also computed the amount of expression variance explained by known technical covariates in each tissue. Based on this, we found that gene expression measurements of whole blood was most confounded, followed by lung, skeletal muscle, tibial nerve, sun-exposed skin, thyroid and subcutaneous adipose. With both WGCNA and graphical lasso, we also found that networks inferred from principal component corrected data were much more sparse compared to uncorrected, and RIN, exonic rate or GC bias corrected counterparts (**Figure 4**). We also observed that tissues that were more confounded, such as blood and lung had denser networks reconstructed by graphical lasso than less confounded tissues such as thyroid (**Figure 4**, **Supplementary Table 2**).

**Figure 4.**
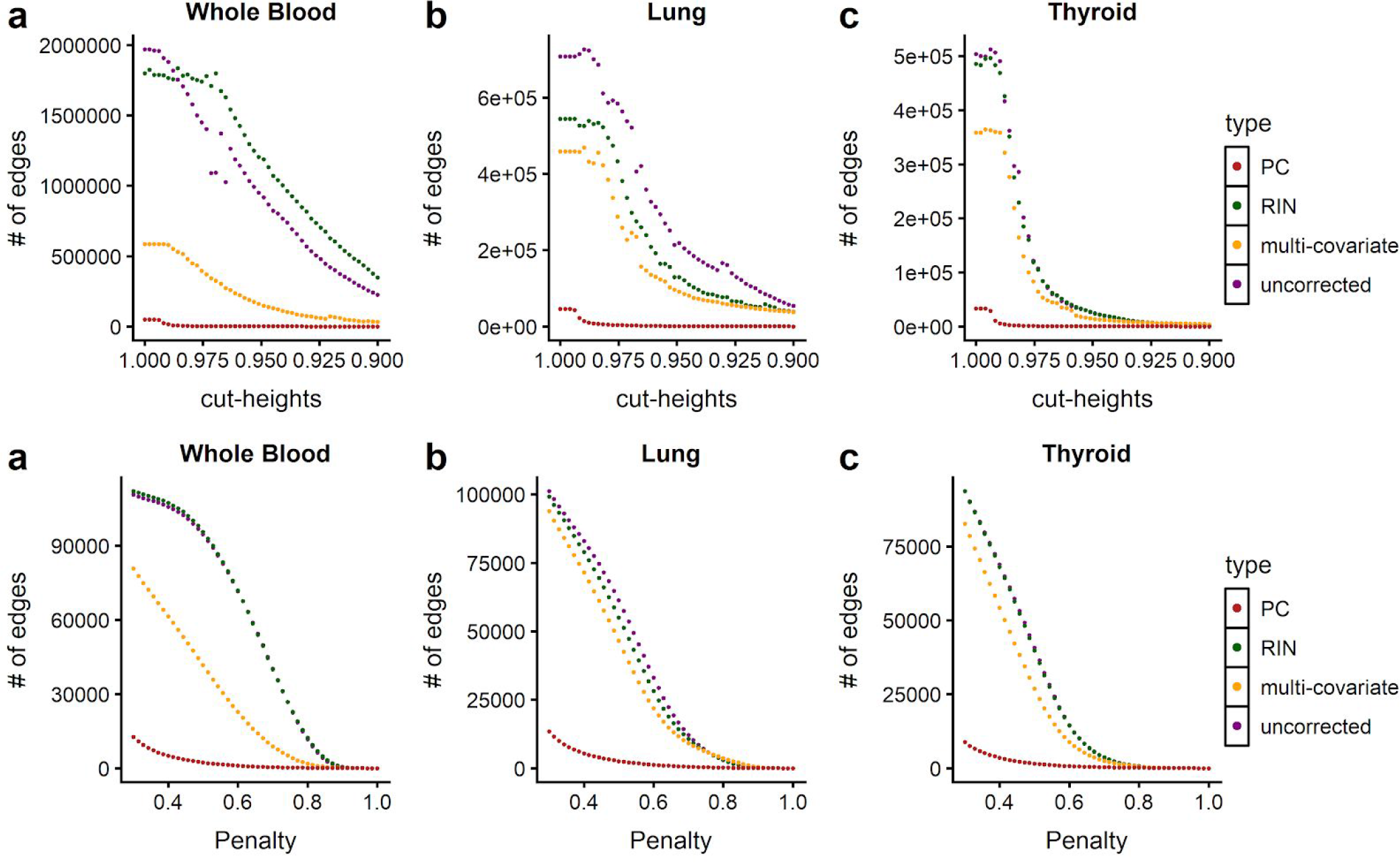
Density of networks inferred from PC-corrected data are sparser. a-c Each point corresponds to number of edges in networks inferred by WGCNA at a cut-height. d-e Each point corresponds to number of edges inferred by graphical lasso in networks obtained at a specific L1 penalty parameter value. Networks inferred by PC corrected data have fewer edges compared to uncorrected or RIN corrected data.

## Discussion and conclusion

Overall we show that network reconstruction methods are vulnerable to latent confounders present in gene expression data. The simulation study demonstrates that graph estimation methods that do not account for confounders make a large number of false discoveries and may lose true dependencies. Similarly, in empirical analysis using GTEx data we see that the networks inferred from the expression data without any correction methods in general performed poorly compared to principal component corrected data. In addition, co-expression networks obtained from gene expression data corrected for effects of RIN, exonic rate, or GC bias individually show little improvement on false discoveries compared to uncorrected data. Thereby our analysis shows that routinely available covariates such as RIN, exonic rate, and GC bias alone are not a sufficient surrogate for the diverse sources of confounding variation in gene expression data and do not fully eliminate patterns of this artifactual variation.. When extensive, well annotated covariates are measured and available, correcting gene expression data jointly with multiple covariates can sometimes improve reconstruction of coexpression networks.

Although, this improvement is still usually not as large as observed for networks reconstructed with PC corrected data. Unfortunately, most studies do not have extensive covariate annotations that could have confounding effects on gene expression and co-expression available. Additionally, a few studies that do have well annotated covariate metadata do not always have them recorded for all samples. Latent factor correction based approaches have been successfully applied to address batch effects and other artifacts in differential expression analysis, GWAS, and eQTL mapping[11,28,29].Our principal component based correction is a simple, yet effective approach to address confounding variation on gene co-expression networks. We do note that overall false discovery rate is high on an absolute scale, regardless of the choice of method and data. However, this is not unique to our analysis, and co-expression network reconstruction frequently yields a high false discovery rate despite enrichment in aggregate for biologically meaningful relationships[6,7].Also, for particularly dense or connected sub-graphs in the underlying biological system that may not match the small-world assumption, removing principal components may remove relevant biological signal and, as with any data cleaning methodology, should be used with caution. We have implemented our PC correction approach in the *sva*Bioconductor package which can be used prior to network reconstruction with a range of methods.

## Methods

### Simulation Example

We constructed a true underlying network with eight nodes that represent genes and three edges that represent conditional dependencies between the genes. Next, we simulated 10,000 observations from a multivariate normal distribution that encode the conditional dependencies corresponding to three edges as non-zero entries in the precision matrix (**Figure 1a**). Then, to introduce confounding in the data, we simulated a sample specific term from a standard normal distribution, and added a scalar multiple of that to genes 2 through 6 (**Figure 1d**). Finally, to correct the data, we regressed out the first principal component from the confounded data (**Figure 1g**). We used graphical lasso to reconstruct networks using the three versions of the data. The code for this simulation example and network reconstruction can be found at: https://github.com/leekgroup/networks_correction/blob/master/publication_rmd/simulation_example_fig1/figure1.Rmd

### Reconstruction of co-expression networks

To evaluate our correction method and its effect on reconstruction of co-expression networks, we used two methods to infer the structure of gene co-expression networks: a) weighted gene co-expression networks (WGCNA)[10] and b) graphical lasso[11]

- WGCNA: Weighted Gene Co-expression Network Analysis (WGCNA) identifies relationships between genes through a power transform of the Pearson correlation coefficient. The algorithm first computes the pairwise correlation between genes *i*and *j*:

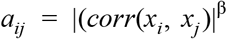

where β > 1. β is the soft-thresholding parameter and it’s value is selected such that the networks obtained are scale free. This is assessed by setting a scale-free topology fit *R*^2^ at least 0.85, between *log*(*p*(*k*)) and *log*(*k*) where *p*(*k*) is the fraction of nodes with at least k neighbors[12, 10, 13, 14].Next, the adjacency matrix between genes is transformed into a topological overlap matrix (TOM) [12]. The TOM is then input into an average linkage hierarchical clustering algorithm to identify network modules - defined as ‘groups of nodes with high topological overlap and indicates high levels of co-expression [12].
- Graphical Lasso: Given our gene expression data contains *N* multivariate gaussian observations each of dimension *p*, i.e. for each observation, we have expression measurements for *p* genes, graphical lasso estimates the structure of the co-expression network over genes by maximizing *L*_1_-penalized log likelihood of a multivariate gaussian with mean μ and covariance Σ over Θ:

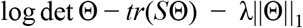 Here *S* is the empirical covariance matrix and Θ = Σ^−1^ is the inverse covariance matrix. The *L*_1_ penalty on Θ induces and controls the amount of sparsity in the solution [11]. Hence, if an entry Θ_*i,j*_ is 0, then variable *i* is conditionally independent of variable *j* given other variables. We used ‘QUIC’ package[15] in R to infer co-expression network structure with graphical lasso.

### Determining sample specific estimates of GC bias

Studies have shown that GC content of genes have significant impact on sequencing read coverage in DNA-seq and RNA-seq experiments. This eventually introduces sample specific biases in expression quantification. To quantify the effect of GC bias, using transcript level fasta files from Gencode v25 we first computed the GC% of each transcript *R* by:

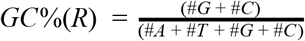

We summarized GC content of genes, by averaging over all transcripts belonging to the gene. Suppose *k* transcripts were transcribed from gene *G*_*i*_ then,

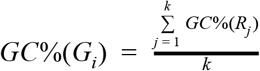

Next using a linear model, we obtain sample specific estimates of GC content of genes:

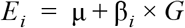

where, *E*_*i*_ is the vector of expression values of all protein coding genes in sample *i*, *G* is the GC content of each gene, and β_*i*_ is the estimate of GC bias in sample *i*.

### Principal component based correction of gene expression

Using a permutation based approach as described in[27], we first determined the number of principal components ‘*p*’to correct the data for with the ‘num.sv’ function in the Bioconductor package *sva*(**Supplementary Table 3**). Next we standardized the data such that the expression of each gene was centered at mean and had unit standard deviation. We compute the principal component loadings *L* of the standardized expression matrix with singular value decomposition (SVD). Using a linear model as described below, we regressed the top ‘*p*’ principal components (*p*as determined by ‘num.sv’) on the each gene *E*_*i*_, from the expression data and computed the residuals *Ê*_*i*_.

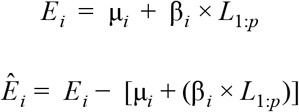

### Reconstruction of co-expression networks with GTEx data

Based on sample size we used gene expression RNAseq data from eight tissues in the GTEx project[14]that included whole blood, lung, skeletal muscle, tibial artery, sun-exposed skin, tibial nerve, subcutaneous adipose, and thyroid. In each tissue we filtered for non-overlapping protein coding genes that had scaled expression (counts scaled by the total coverage of the sample) of at least 0.1 ≥ 25% of total number of observations. Next, we log2 transformed the scaled gene expression data, and performed following steps to select the most variable 5000 genes across all tissues.

- For each tissue, assign a rank to each gene by variance, such that the most variable gene is ranked first and least variable gene is ranked in last.
- Using the ranked list of genes from each tissue, assign an average rank to each gene across eight tissues.
- Select top 5000 genes from the average rank list for network reconstruction.

We used multiple approaches to correct gene expression data from each tissue individually as described below:

- Residuals from RIN/Exonic Rate/ GC bias: Using a linear model, we regressed the RNA integrity number (RIN), exonic rate or sample specific estimate of GC bias on the expression data and computed the residuals
- Residuals from multiple covariate correction: In each tissue individually, we estimated expression percent variance *R*^2^ explained by the known technical confounders. Next, using a linear model we regressed the technical covariates with *R*^2^ ≥ 0.01 in a tissue and computed the residuals. **(Supplementary Table 4**)
- Residuals from principal components: For each tissue, principal component based gene expression residuals were computed as described in above.

Prior to reconstructing co-expression networks with WGCNA and graphical lasso, we transformed the uncorrected and corrected expression of each gene to a Gaussian distribution by projecting the expression of each gene to the quantiles of a standard normal.

To reconstruct signed weighted gene co-expression networks with WGCNA, we identified lowest power for which scale-free fit *R*^2^ between *log*(*p*(*k*)) and *log*(*k*) exceeds 0.85. Here *p*(*k*) is the fraction of nodes in the network with at least *k* neighbors. Then we used the *‘blockwisemodules’*function in the WGCNA CRAN package to identify co-expression modules at varying cut-heights of hierarchical dendrogram ranging from 0.9 to 1.0. For networks reconstructed with WGCNA, we considered all genes in the same module to be a fully-connected subgraph.

For reconstruction of co-expression networks with graphical lasso, we first computed the gene covariance matrix and used then used ‵QUIC‵ function in the *QUIC* R package to infer co-expression networks with penalization parameter λ ranging from 0.3 to 1.0

### Evaluation of co-expression networks

Since the underlying network structure is generally unknown, we used genes known to be functional in the same pathways as ground truth to assess these networks.

- Canonical pathway databases: We downloaded the latest pathway information (2016) from KEGG, Biocarta and Pathway Interaction Database from Enrichr[18, 19], that were also annotated as canonical pathways by MSigDB[37]. The number of pathways/genesets in each of these databases were:

– KEGG - 293
– Biocarta - 237
– Reactome - 1530
– Pathway Interaction Database - 209 Any pair of genes that have at least one pathway in common were assumed as true connection. An edge that was observed between a pair of genes in the inferred network (from WGCNA or graphical lasso) and was also present in the list of real connections was called as a true positive (TP). We defined false positive (FP) to be an edge that was observed between a pair of genes in the inferred network, however was absent in the list of real connections. False negatives (FN) were the edges that were missing in the inferred network but were present in the list of real connections. Using this definition of true positive, false positive and false negative, we compute false discovery rate for the networks inferred by WGCNA and graphical lasso with different forms of corrected and uncorrected data.
- Shared true positives: We obtained a refined list of real connections described above by restricting to pairs of genes that were present in at least two pathway databases.

All TP, FP and FN were computed with genes restricted to the most variable 5000 genes that were used for reconstructing co-expression networks. Using the above mentioned definitions of TP, FP and FN, we compute false discovery rate as given below:

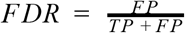

## List of Abbreviations

WGCNA: Weighted gene co-expression networks

## Declarations

### Ethics approval and consent to participate

**Consent for publication**

## Availability of data and materials

All analyses was performed using R and scripts are available on github at: https://github.com/leekgroup/networks_correction

